# Lineage Detector: Efficient Tool for Detecting New SARS-Cov-2 Lineages

**DOI:** 10.1101/2024.11.01.621557

**Authors:** Xu Zou

## Abstract

Since the novel virus of SARS-Cov-2’s emergence, it continues to mutate at significant speed. The mutation speed and diversity of the virus has reached a level that is hard to be analysed purely via human tracing even with the help of UShER trees.

We create an open-sourced tool of *Lineage Detector*, an automated tool that helps highlight the most important lineages of interest on Usher trees. *Lineage Detector* can highlight the most interesting SARS-CoV-2 branches for manual investigation and reduce the work of volunteer variant hunters to 8 ∼ 10%, greatly improve their efficiency.

Since its release, it has helped the identification, proposal and designation process of more than 100 SARS-CoV-2 variants.

## 1 INTRODUCTION

Since the novel coronavirus of SARS-Cov-2 being introduced to human population, it starts evolving at extraordinary speed, especially after all governments give up attempts to control it [2, 10].

The fast evolution of SARS-Cov-2 posts a great challenge on variant tracking and identification. Pango [6] serves as an authoritative classification system of SARS-Cov-2 variants. Pango lineages are designated only if they contain a sufficient number of sequences with high genome coverage with some significant mutations [6]. Up to October 2024, there are already as many as 4263 different SARS-Cov-2 lineages identified by Pango.

Such many of the lineage are mainly identified by the automated evolutionary tree placement tool of UShER (Ultrafast Sample placement on Existing tRee) [9]. The tool collects genetic samples of SARS-Cov-2 from various sources to build a phylogenetic tree that clarifies SARS-Cov-2 variants. New lineages are usually idenfied via manually looking at a sub-branch of the UShER tree.

The identification process of a new SARS-Cov-2 lineage involves 2 stages:

- Firstly, volunteers submit sequences of SARS-Cov-2 to UShER to see their placements. If some of the seqs belong to an interesting branch, that they gather together on a sub-branch of some existing variants and shares some unique mutations of interest, volunteers may make a proposal on the pango proposal repository ^1^.
- Secondly, Pango officials will verify each proposal and decide whether to designate it as a new variant.

As mutations become more and more complex, it becomes increasingly difficult to find all of the new lineages manually by looking at different sub-branches of the UShER tree. Hence we propose *Lineage Detector*, an automated tool to highlight important new lineages on UShER tree. *Lineage Detector* can highlight the most important 8 ∼ 10% of the sequences from the mass GISAID dataset for manual investigation. It has helped identification of more than 100 SARS-Cov-2 variants.

## 2 METHODOLOGY

### 2.1 The UShER Framework

Currently, UShER serves as the major tool for SARS-Cov-2 variant classification and identification. It is an algorithm of putting new sequences based on an existing phylogenetic tree. Given a phylogenetic tree and a new sequence, UShER finds the closest place on the tree to the sequence and create a branch for the sequence.

Figure 2 illustrates how usher places new sequences on an existing phylogenetic tree. It uses intermediate nodes to support the formation of an evolution tree. For new sequences, it finds a position that the new sequence needs the fewest number of mutations to insert in, and add intermediate nodes to the tree if necessary.

**Figure 1:**
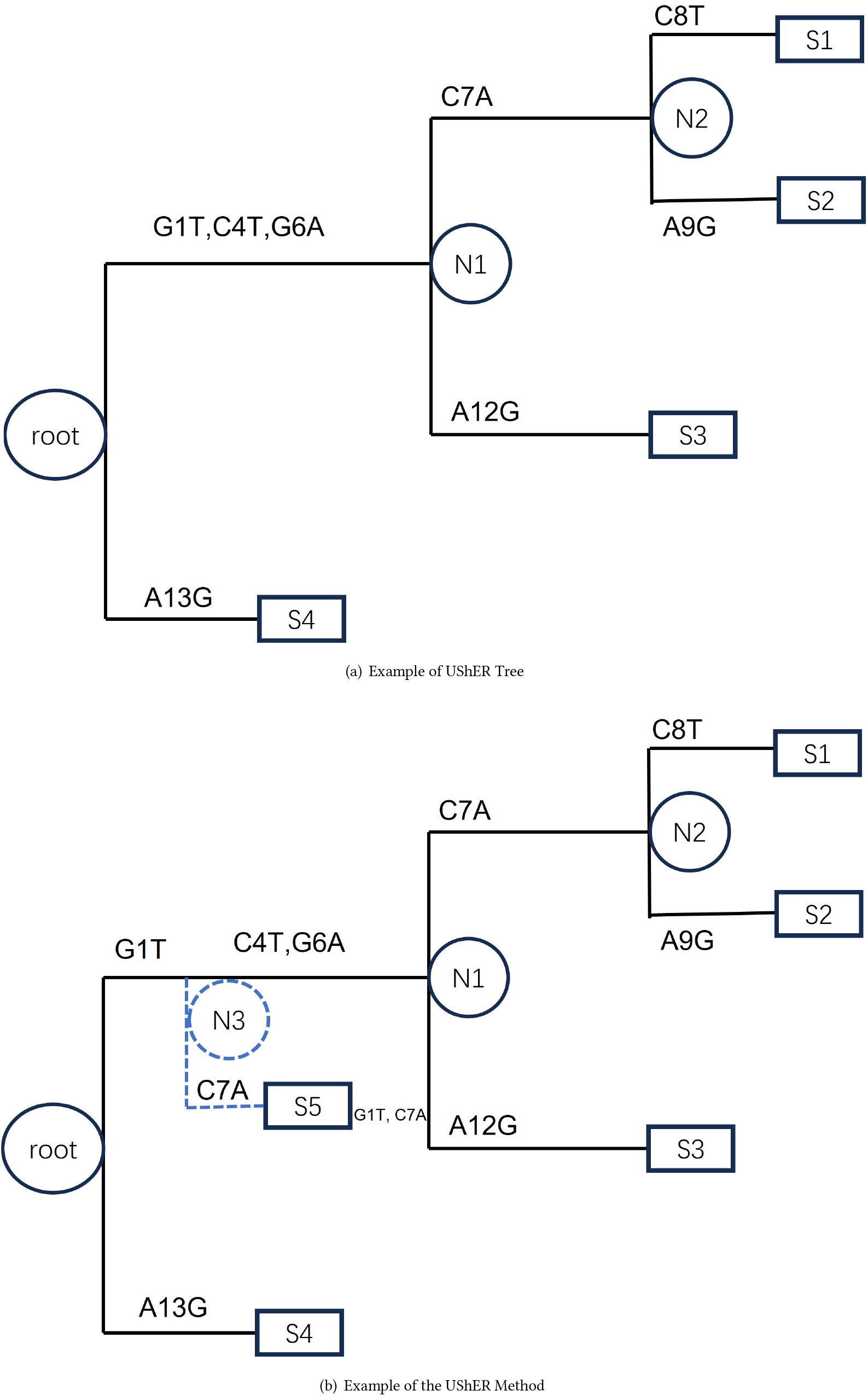
Example of how UShER places new sequences on an existing tree. It uses intermediate nodes to support the formation of an evolution tree. It finds a position that the new sequence needs the minimal number of mutations to insert in, and add intermediate nodes to the tree if necessary.

**Figure 2:**
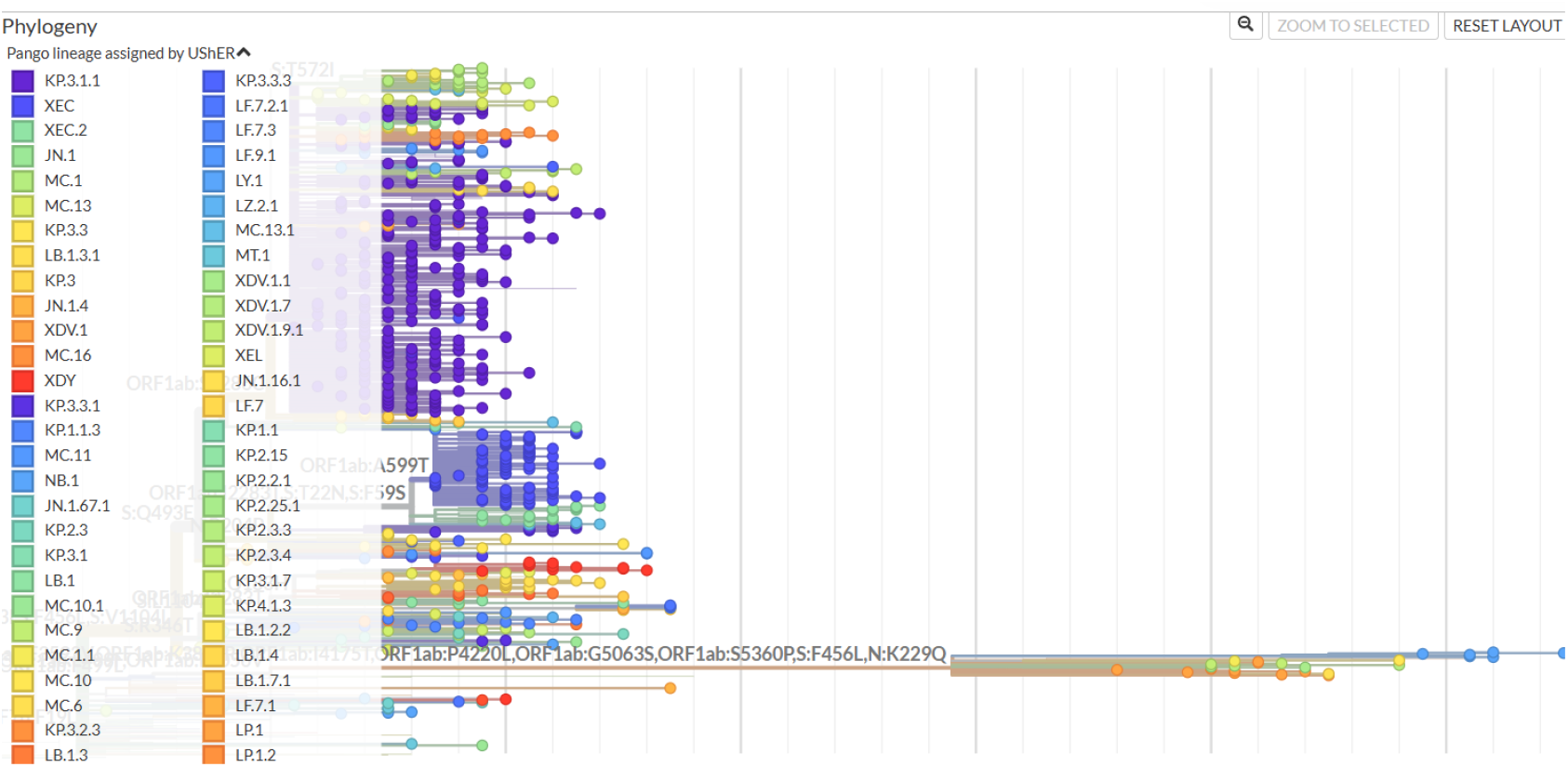
An example of UShER tree classification for the most recent 1,000 sequences(by sampling date) that are uploaded from 2024-10-23 to 2024-10-26.

Genetic sequences may not be 100% accurate. They may have long part of missing coverage, or sequencing artefacts that wrongly sequence some of the codons. Hence UShER apply many manual modifications, it blocks sites that are likely to cause artefacts on specific branches and it removes sequences that are believed not being sequenced correctly. These manual modifications are rapidly updated according to the most recent data and community suggestions. These manual maintainence works make UShER avoid most of the errors in tree building and becomes the best source of SARS-CoV-2 phylogenetic tree.

### 2.2 Non-overlapping and Overlapping Genes

There exists various open reading frames on SARS-Cov-2. An open reading frame starts with a start codon, and keeps translating nucleocides to amino acids until a stop codon. Reading from 5’ start to 3’ end, starts translation at a start codon and stops at a stop codon. The translated genes are called non-overlapping gene. They don’t overlap with each other and an ordered translation will yield these genes. UShER only displays mutations on non-overlapping genes.

On the other hand, overlapping genes refers to open reading frames that overlaps with other open reading frames. These genes have their start codons placed at a place where a non-overlapping gene is translating. These genes won’t be translated if the translation process is fully in order. However, under some conditions these genes may be translated and result in real effects. The most famous example is the Orf9b gene in SARS-Cov-2. It is an overlapping gene overlaps with the N protein. It has functions on immune response of SARS-CoV-2 infections. [3, 5] Notably, T28297C, a synonymous mutation on N protein, adds great fitness to XBB in 2023, and forms two main SARS-CoV-2 clades, EG.5.1 and XBB.1.16. This fitness gain is believed to come from a non-synonymous Orf9b mutation, Orf9b:I5T, highlighting the importance of mutations on overlapping genes in SARS-CoV-2 evolution. [4, 8]

### 2.3 Recombinants

Recombination is a common evolutionary tool for RNA viruses. SARS-CoV-2 variants may recombine with each other and create recombinants. [7] Recombinants take part of variant A and part of variant B, hence they may behave like having a lot of reversions from either A or B, which may trigger then data-cleaning filter of UShER that removes sequences it believe to have more than 5 reversions due to low sequencing quality.

### 2.4 Pango Designation and Nextclade Clade

Currently, SARS-CoV-2 variants are mainly traced by the Pango project [6]. It gives every lineage with notable mutations a unique label. The system starts from *A* and *B*, and denotes descendants of variants by adding .*x*. When a descendant reaches a level of 3, a new letter starting from *C* is picked to simplify the naming. *X* is used to label recombinants. Starting from *XA*, designated recombinants pick up a unique letter in alphabetical order and adds an *X* before it. Currently, the most recently designated variant is *N M*.1, which is a simplified form of *KP* .2.3.4.1, or *J N* .1.11.1.2.3.4.1, or *BA*.2.86.1.1.11.1.2.3.4.1, or *B*.1.1.529.2.86.1.1.11.1.2.3.4.1, and *X EN*, which is a recombinant of *KP* .1.1.6 and *J N* .1.11.1.

Most of the notable mutations are on spike. As illustrated in Table 1, more than half of the designated non-recombinant variants adds at least one spike mutation on top of their parents.

**Table 1:**
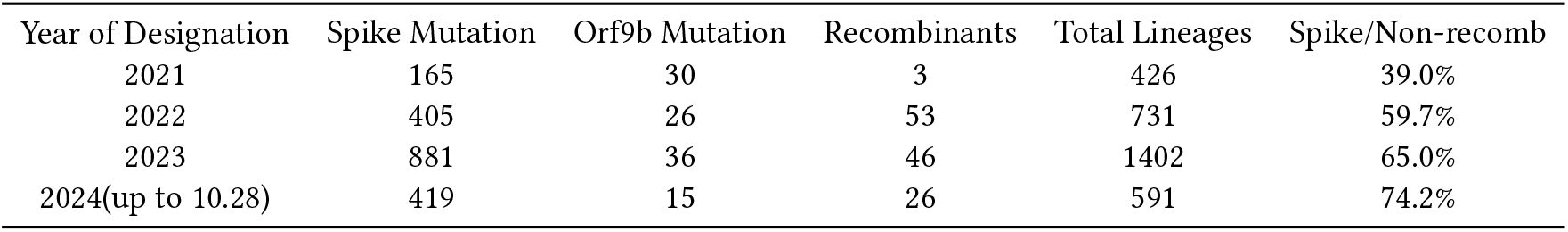
Defining mutations of UShER Linages.

Nextclade [1] is a more coarse grain variant categorization method. Only variants that both mutates and spreads significantly, either getting a WHO naming, or become continental or global dominant at some time point and with significant spike mutations, are designated a nextclade clade. As WHO has stopped naming new SARS-CoV-2 variants since 2021, only variants that have significant spike mutations and achieves or to achieve continental domination(30% prevalence or 5% prevalence when daily growth is above 5%) or global domination(20% prevalence) can be lifted to nextclade clades. These variants are major variants that have significant effects on the level of the pandemic. Table 2 displays number of nextclade clades each year.

**Table 2:** Number of Nextclade Clades Each Year.

| Year | 2019 | 2020 | 2021 | 2022 | 2023 | 2024(up to 10-31) |
| --- | --- | --- | --- | --- | --- | --- |
| Nextclade Clades via WHO Naming | 0 | 2 | 8 | 0 | 0 | 0 |
| Nextclade Clades via Intrinsic Significance | 2 | 8 | 5 | 6 | 9 | 7 |

### 2.5 Lineage Detector

Every day, thousands of new sequences are added to GISAID, volunteers(variant hunters) analyse them using UShER and propose potential new lineages to Pango. However, due to having to look at too many sequences, this workload is heavy and easy to miss.

Figure 2 illustrates an example of an UShER tree classification result when submitting the ost recent 1,000 sequences uploaded from 2024-10-23 to 2024-10-26. As can be seen, the resulting tree is very messy and includes hundreds of separate branches.

Hence we develop *Lineage Detector*, it is a tool that highlights important lineages on usher trees that deserve proposals automatically. With the implementation of *Lineage Detector*, workload of variant hunters can be largely reduced.

Figure 3 shows the same tree of figure 2 with hightlighted branches and samples. It reduces variant hunters’ workload from wandering within hundreds of lineages to only an XDV.1 branch, a KP.3.3 branch, an XEC.2 branch and several KP.3.1.1 branches, significantly improves their efficiency.

**Figure 3:**
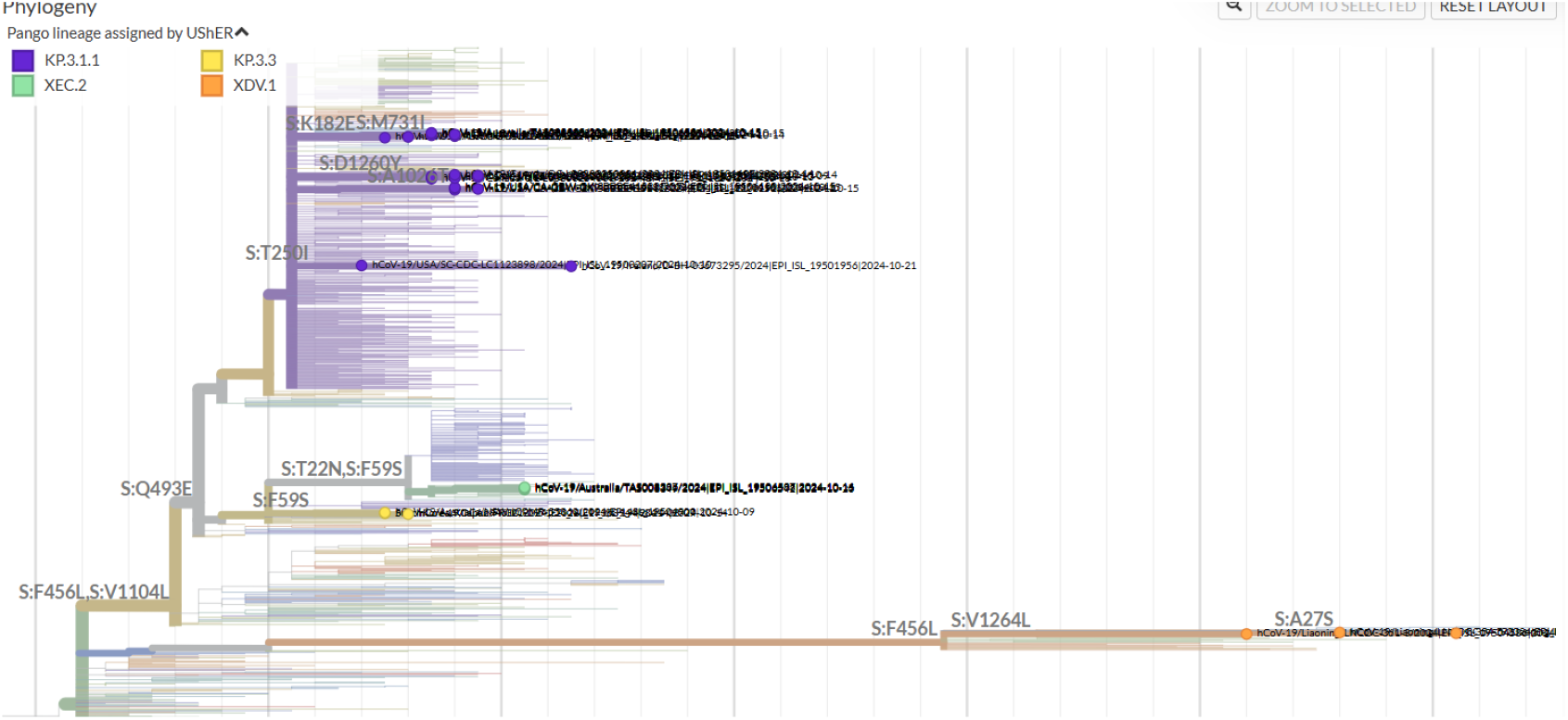
UShER tree classification for the most recent 1,000 sequences(by sampling date) that are uploaded from 2024-10-23 to 2024-10-26, sequences deserve attention are highlighted.

Based on above analyse in section **??**, *Lineage Detector* focuses on highlighting the following lineages.

- Lineages with additional spike mutations from a undesignated lineage. The new spike mutations may grant the lineage additional fitness via transmission rate change or immune escape, and help it to spread.
- Lineages with additional Orf9b mutations from a undesignated lineage. As an overlapping gene, Orf9b mutations are not present on the UShER tree. However, we can detect them by analysing the nucleocide mutations, and highlight them so that we won’t miss these mutations.
- Lineages with additional stop codon or removal of start codon of some proteins. These mutations may change the virus at large scale and deserves high-light.
- Sequences with more than 5 reversions after a designated lineage. These sequences may be automatically removed in future UShER trees, and they deserve an investigation to see whether they are actually recombinants so as to manually add them back to the UShER tree.

Based on the above criteria, *Lineage Detector* is published as an open-sourced github repository ^2^ to highlight important lineages for variant hunters.

## 3 IMPLEMENTATION

### 3.1 Maintainence of Recent High-lighted Samples

*Lineage Detector* updates trees of the most recent samples every a few days, finds and highlights important lineages rapidly.

Every few days, new samples from GISAID are downloaded, they are separated to batches of less than 1,000 according to their collection dates. Each batch is then submitted to UShER to analyse their placements. The results are further analysed by *Lineage Detector* to highlight important lineages deserve manual attention according to criteria described in section 2.5. The resulting files are publicly available and can be viewed easily using nextclade [1]. An example way to view the analysed tree is through link https://nextstrain.org/fetch/raw.githubusercontent.com/xz-keg/Lineage-Detector/main/daterank.json?f_userOrOld=highlighted%20sample, where date refers to the time analysed and is in the form YYYY-MM-DD, rank refers to the rank of the batch on that day.

For important lineages, we require the lineage to have at sequences from 2 different places in a batch to be high-lighted, or have at least 3 sequences. Sequences from some countries known for submitting clusters count as 0.5.

We also open-source the code so as to allow personal analyse on any batches of sequences with flexible thresholds.

### 3.2 Results

*Lineage Detector* highlights new SARS-CoV-2 lineages that may be important for variant hunters, greatly improving their efficiency.

Table 3 takes July, August, September and October as examples and show the number of highlighted sequences in these months compared to total number of sequences. As can be seen, *Lineage Detector* highlights only 8.85% of the total sequences analysed, greatly reducing the workload of variant hunters. Despite sequences highlighted being only a small proportion of total sequences, these sequences are really the important ones. Since the tool is publicized, many novel lineages are found with the help of it. Table 4 lists designated SARS-CoV-2 variants found with the help of *Lineage Detector*. Up to 2024-10-30, *Lineage Detector* has helped finding 109 novel variants, including KP.2.3 and XEC which grows significantly to be lifted to nextclade clade, six other recombinants and 101 other variants.

**Table 3:** Number of Highlighted Sequences Compared with Total Sequences in July, August, September and October 2024.

| Time | Highlighted Sequences | Total Sequences Analysed | Highlight Rate |
| --- | --- | --- | --- |
| 2024-7 | 6411 | 59333 | 10.81% |
| 2024-8 | 6319 | 74600 | 8.47% |
| 2024-9 | 4968 | 61257 | 8.11% |
| 2024-10 | 4417 | 54693 | 8.08% |
| Total | 22115 | 249883 | 8.85% |

**Table 4:** List of Pango lineages found and proposed involving the help of *Lineage Detector* up to 2024-10-30.

| Pango Lineages Lifted to Nextclade Clade |  |  |  |  |  |  |  |
| --- | --- | --- | --- | --- | --- | --- | --- |
| KP.2.3(24G) |  |  |  | XEC(24F) |  |  |  |
| Other Recombinant Lineages |  |  |  |  |  |  |  |
| XDW | XED | XEE | XEL | XEF | XEJ |  |  |
| Other Pango Lineages |  |  |  |  |  |  |  |
| LF.1.1 | LF.1.1.1 | XDD.1.1.1 | JN.1.40 | JN.1.57.1 | XDK.4.1 | JN.1.42.1 | XDK.1.1 |
| JN.1.4.8 | LF.2 | KP.1.1.3 | KP.2.19 | KP.2.22 | KP.2.23 | KP.2.24 | KP.4.2.4 |
| XDQ.3 | KP.1.1.6 | XDK.3 | JN.1.63 | JN.1.63.1 | XDQ.1.1 | KP.2.4 | KP.3.3 |
| KP.1.1.2 | KR.1.1 | KP.2.8 | KP.2.10 | KP.3.2.1 | XDV.1.3 | MK.1 | MK.2 |
| XDV.1.2 | XDV.1.2.1 | LP.1.1 | LB.1.7.1 | KP.2.2.2 | KP.2.3.5 | KP.3.3.1 | LU.2 |
| KP.4.2.5 | KP.6 | MN.1 | MD.3.1.4 | MC.2 | MC.2.1 | KP.3.1.7 | MM.2 |
| LB.1.3.3 | LF.9 | LF.9.1 | LD.2 | KP.2.20.1 | KP.2.3.8 | KP.2.3.9 | NG.1 |
| KS.1.4 | LP.6 | KP.2.25.2 | MC.4 | LW.1.1 | MU.2.1 | MC.8.1 | MC.5 |
| KP.2.15.1 | LB.1.2.2 | MC.14 | MQ.2 | XDV.1.5 | NB.1 | KP.2.3.12 | MC.11.1 |
| LF.8 | LF.8.1 | LF.8.1.1 | MC.12 | KP.3.2.6 | LZ.2.1.2 | ND.2 | ND.1.1 |
| MC.10.1 | XDV.1.9.1 | MC.17 | NC.1 | LF.7.4 | LZ.2.1.1 | NA.1.1 | JN.1.59.1 |
| NK.1 | LF.7.1.2 | LF.7.2.1 | NJ.1 | KP.3.1.9 | LU.2.2 | KP.3.1.10 | LB.1.4.3 |
| MC.1.1 | MD.3.1.3 | NM.1 | ND.3 | NE.1 |  |  |  |

## 4 CONCLUSION

In this report, we propose *Lineage Detector*, a useful automated lineage detection tool that help filtering the most important 8 ∼ 10% lineages from mass SARS-CoV-2 sequence dataset for manual investigation. Since its release, it has helped variant hunters to find and propose more than 100 SARS-CoV-2 Pango variants, including two major variants that were lifted to nextclade clades subsequently.

## Footnotes

1 https://github.com/cov-lineages/pango-designation

2 https://github.com/xz-keg/Lineage-Detector

## REFERENCES

[1] Ivan Aksamentov, Cornelius Roemer, Emma B Hodcroft, and Richard A Neher. Nextclade: clade assignment, mutation calling and quality control for viral genomes. Journal of open source software, 6(67):3773, 2021.

[2] Carlos M Duarte, David I Ketcheson, Víctor M Eguíluz, Susana Agustí, Juan Fernández-Gracia, Tahira Jamil, Elisa Laiolo, Takashi Gojobori, and Intikhab Alam. Rapid evolution of sars-cov-2 challenges human defenses. Scientific Reports, 12(1):6457, 2022.

[3] He-wei Jiang, Hai-nan Zhang, Qing-feng Meng, Jia Xie, Yang Li, Hong Chen, Yun-xiao Zheng, Xue-ning Wang, Huan Qi, Jing Zhang, et al. Sars-cov-2 orf9b suppresses type i interferon responses by targeting tom70. Cellular & molecular immunology, 17(9):998–1000, 2020.

[4] Rajesh P Karyakarte, Rashmita Das, Mansi V Rajmane, Sonali Dudhate, Jeanne Agarasen, Praveena Pillai, Priyanka M Chandankhede, Rutika S Labhshetwar, Yogita Gadiyal, Preeti P Kulkarni, et al. Chasing sars-cov-2 xbb. 1.16 recombinant lineage in india and the clinical profile of xbb. 1.16 cases in maharashtra, india. Cureus, 15(6), 2023.

[5] Svenja Lenhard, Sarah Gerlich, Azkia Khan, Saskia Rödl, Jan-Eric Bökenkamp, Esra Peker, Christine Zarges, Janina Faust, Zuzana Storchova, Markus Räschle, et al. The orf9b protein of sars-cov-2 modulates mitochondrial protein biogenesis. Journal of Cell Biology, 222(10):e202303002, 2023.

[6] Áine O’Toole, Oliver G Pybus, Michael E Abram, Elizabeth J Kelly, and Andrew Rambaut. Pango lineage designation and assignment using sars-cov-2 spike gene nucleotide sequences. BMC genomics, 23(1):121, 2022.

[7] Orsolya Anna Pipek, Anna Medgyes-Horváth, József Stéger, Krisztián Papp, Dávid Visontai, Marion Koopmans, David Nieuwenhuijse, Bas B Oude Munnink, VEO Technical Working Group Cochrane Guy 3 Rahman Nadim 3 Cummins Carla 3 Yuan David Yu 3 Selvakumar Sandeep 3 Mansurova Milena 3 O’Cathail Colman 3 Sokolov Alexey 3 Thorne Ross 3 Worp Nathalie 2 Amid Clara 2, and István Csabai. Systematic detection of co-infection and intra-host recombination in more than 2 million global sars-cov-2 samples. Nature Communications, 15(1):517, 2024.

[8] Shuhei Tsujino, Sayaka Deguchi, Tomo Nomai, Miguel Padilla-Blanco, Arnon Plianchaisuk, Lei Wang, MST Monira Begum, Keiya Uriu, Keita Mizuma, Naganori Nao, et al. Virological characteristics of the sars-cov-2 omicron eg. 5.1 variant. Microbiology and immunology, 68(9):305–330, 2024.

[9] Yatish Turakhia, Bryan Thornlow, Angie S Hinrichs, Nicola De Maio, Landen Gozashti, Robert Lanfear, David Haussler, and Russell Corbett-Detig. Ultrafast sample placement on existing trees (usher) enables real-time phylogenetics for the sars-cov-2 pandemic. Nature genetics, 53(6):809–816, 2021.

[10] Sijie Yang, Yuanling Yu, Yanli Xu, Fanchong Jian, Weiliang Song, Ayijiang Yisimayi, Peng Wang, Jing Wang, Jingyi Liu, Lingling Yu, et al. Fast evolution of sars-cov-2 ba. 2.86 to jn. 1 under heavy immune pressure. The Lancet Infectious Diseases, 24(2):e70–e72, 2024.

